# Allele-specific genome editing using CRISPR-Cas9 causes off-target mutations in diploid yeast

**DOI:** 10.1101/397984

**Authors:** Arthur R. Gorter de Vries, Lucas G. F. Couwenberg, Marcel van den Broek, Pilar de la Torre Cortés, Jolanda ter Horst, Jack T. Pronk, Jean-Marc G. Daran

## Abstract

Targeted DNA double-strand breaks (DSBs) with CRISPR-Cas9 have revolutionized genetic modification by enabling efficient genome editing in a broad range of eukaryotic systems. Accurate gene editing is possible with near-perfect efficiency in haploid or (predominantly) homozygous genomes. However, genomes exhibiting polyploidy and/or high degrees of heterozygosity are less amenable to genetic modification. Here, we report an up to 99-fold lower gene editing efficiency when editing individual heterozygous loci in the yeast genome. Moreover, Cas9-mediated introduction of a DSB resulted in large scale loss of heterozygosity affecting DNA regions up to 360 kb that resulted in introduction of nearly 1700 off-target mutations, due to replacement of sequences on the targeted chromosome by corresponding sequences from its non-targeted homolog. The observed patterns of loss of heterozygosity were consistent with homology directed repair. The extent and frequency of loss of heterozygosity represent a novel mutagenic side-effect of Cas9-mediated genome editing, which would have to be taken into account in eukaryotic gene editing. In addition to contributing to the limited genetic amenability of heterozygous yeasts, Cas9-mediated loss of heterozygosity could be particularly deleterious for human gene therapy, as loss of heterozygous functional copies of anti-proliferative and pro-apoptotic genes is a known path to cancer.

## INTRODUCTION

CRISPR-Cas9-assisted genome editing requires the simultaneous presence of the Cas9 endonuclease and a guide-RNA (gRNA) that confers target-sequence specificity (1). A gRNA consists of a structural domain and a variable sequence homologous to the targeted sequence (1-4). A Cas9-gRNA complex introduces a DSB when the gRNA binds to its reverse complement sequence on the 5’ side of a PAM sequence (NGG). Imperfect gRNA complementarity and/or absence of a PAM sequence strongly reduce editing efficiencies (5). CRISPR-Cas9 enables specific editing of any sequence proximal to a PAM sequence, with minimal off-targeting effects (5). The introduction of a DSB facilitates genome editing by increasing the rate of repair by homologous recombination (6). When a repair fragment consisting of a DNA oligomer with homology to regions on both sides of the introduced DSB is added, it is integrated at the targeted locus by homologous recombination, resulting in replacement of the original sequence and repair of the DSB (2-4). In *S. cerevisiae*, double stranded DNA oligomers with 60 bp of homology are sufficient to obtain accurate gene-editing in almost 100% of transformed cells (3). By inserting sequences between the homologous regions of the repair oligonucleotide, heterozygous sequences of up to 35 Kbp could be inserted at targeted loci (7). While such gene editing approaches have been very efficient in haploid and homozygous diploid yeasts, the accurate introduction of short DNA fragments can be tedious in heterozygous yeast. In homozygous diploid and polyploid eukaryotes, CRISPR-Cas9 introduces DSBs in all alleles of a targeted sequence (8). In heterozygous genomes, gRNAs can be designed for allele-specific targeting if heterozygous loci have different PAM motifs and/or different 5’ sequences close to a PAM motif (8,9), enabling allele-specific gene editing using Cas9. In such cases, a DSB is introduced in only one of the homologous chromosomes while the other homolog remains intact. However, the presence of intact homologous chromosomes facilitates repair of DSBs by homologous recombination (HR), homology-directed repair (HDR) or break-induced repair (BIR) in eukaryotes (10-12). Therefore, the presence of an intact homologous chromosome could provide an alternative path of DSB repair and, thereby, compete with the intended gene-editing event. So far, no systematic analysis has been performed of the efficiency of Cas-9-mediated gene editing at heterozygous loci. To investigate if Cas9 gene editing works differently in heterozygous diploid yeast, we tested if allele-specific targeting of heterozygous loci using Cas9 enables accurate gene editing in an interspecies *Saccharomyces* hybrid, and investigated the resulting transformants. In addition, we systematically investigated the efficiency of CRISPR-Cas9-mediated genome editing when targeting various homozygous and heterozygous loci in diploid laboratory *Saccharomyces cerevisiae* strains while monitoring off-target mutations.

## MATERIAL AND METHODS

### Strains, plasmids, primers and statistical analysis

*S. cerevisiae* strains used in this study are derived from the laboratory strains CEN.PK113-7D and S288C (13,14). Yeast strains, plasmids and oligonucleotide primers used in this study are provided in Tables S3, S4 and S5. Statistical significance was determined using two-tailed unpaired Student’s t-tests in GraphPad Prism 4.

### Media and growth conditions

Plasmids were propagated overnight in *Escherichia coli* XL1-Blue cells in 10 mL LB medium containing 10 g·L^-1^ peptone, 5 g·L^-1^ Bacto Yeast extract, 5 g·L^-1^ NaCl and 100 mg·L^-1^ ampicillin at 37°C. Unless indicated otherwise, yeast strains were grown at 30 °C and 200 RPM in 100 mL shake flasks containing 50 mL YPD medium, containing 10 g·L^-1^ Bacto yeast extract, 20 g·L^-1^ Bacto peptone, and 20 g·L^-1^ glucose. Alternatively, strains were grown in synthetic medium (SM) containing 6.6 g·L^-1^ K_2_SO_4_, 3.0 g·L^-1^ KH_2_PO_4_, 0.5 g·L^-1^ MgSO_4_·7H_2_O, 1 mL·L^-1^ trace elements, 1 mL·L^-1^ vitamin solution (15) and 20 g·L^-1^ glucose. For uracil auxotrophic strains, SM-derived media were supplemented with 150 mg·L^-1^ uracil (16). Solid media were supplemented with 20 g·L^-1^ agar. Selection for the amdS marker was performed on SM-AC: SM medium with 0.6 g·L^-1^ acetamide as nitrogen source instead of (NH_4_)_2_SO_4_ (17). The amdS marker was lost by growth on YPD and counter-selected on SM-FAC: SM supplemented with 2.3 g·L^-1^ fluoroacetamide (17). Yeast strains and *E. coli* containing plasmids were stocked in 1 mL aliquots after addition of 30% v/v glycerol to the cultures and stored at -80 °C.

### Flow cytometric analysis

Overnight aerobic cultures in 100 mL shake flasks on 20 mL YPD medium were vortexed thoroughly to disrupt cell aggregates and used for flow cytometry on a BD FACSAria™ II SORP Cell Sorter (BD Biosciences, Franklin Lakes, NJ, USA) equipped with 355 nm, 445 nm, 488 nm, 561 nm and 640 nm lasers and a 70 µm nozzle, and operated with filtered FACSFlow™ (BD Biosciences). Cytometer performance was evaluated prior to each experiment by running a CST cycle with CS&T Beads (BD Biosciences). Drop delay for sorting was determined by running an Auto Drop Delay cycle with Accudrop Beads (BD Biosciences). Cell morphology was analysed by plotting forward scatter (FSC) against side scatter (SSC). The fluorophore mRuby2 was excited by the 561 nm laser and emission was detected through a 582 nm bandpass filter with a bandwidth of 15 nm. The fluorophore mTurquoise2 was excited by the 445 nm laser and emission was detected through a 525 nm bandpass filter with a bandwidth of 50 nm. The fluorophore Venus was excited by the 488 nm laser and emission was detected through a 545 nm bandpass filter with a bandwidth of 30 nm. For each sample, 100’000 events were analysed and the same gating strategy was applied to all samples of the same strain. First, “doublet” events were discarded on a FSC-A/FSC-H plot, resulting in at least 75’000 single cells for each sample. Of the remaining single cells, cells with and cells without fluorescence from Venus were selected in a FSC-A/Venus plot. For both these groups, cells positive for mRuby2 and mTurquoise2, cells positive for only mRuby2, cells positive for only mTurquoise2 and cells negative for mRuby2 and mTurquoise2 were gated. The same gating was used for all samples of each strain. Sorting regions (‘gates’) were set on these plots to determine the types of cells to be sorted. Gated single cells were sorted in 96-well microtiter plates containing YPD using a “single cell” sorting mask, corresponding to a yield mask of 0, a purity mask of 32 and a phase mask of 16. FACS data was analysed using FlowJo^®^ software (version 3.05230, FlowJo, LLC, Ashland, OR). Separate gating strategies were made for IMX1555, IMX1557 and IMX1585 to account for possible differences in cell size, shape and morphology.

### Plasmid assembly

Plasmid pUD574 was de novo synthesised at GeneArt (Thermo Fisher Scientific, Waltham, MA) containing the sequence 5’ GGTCTCGCAAAATTACACTGATGAGTCCGTGAGGACGAAACGAGTAAGCTCGTCTGTAATATCTTAATGCTAAAGTTTTAGAGCTAGAAATAGCAAGTTAAAATAAGGCTAGTCCGTTATCAACTTGAAAAAGTGGCACCGAGTCGGTGCTTTTGGCCGGCATGGTCCCAGCCTCCTCGCTGGCGCCGGCTGGGCAACATGCTTCGGCATGGCGAATGGGACACAGCGAGACC 3’.

Plasmids pUD429 was constructed in a 10 µL golden gate assembly using T4 ligase (St. Louis, MO, USA) and BsaI (New England BioLabs, Ipswich, MA) from 10 ng of parts pYTK002, pYTK047, pYTK067, pYTK079, pYTK081 and pYTK083 of the yeast toolkit as described previously (18). Similarly, pUD430 was constructed from pYTK003, pYTK047, pYTK068, pYTK079, pYTK081 and pYTK083, and pUDP431 from pYTK004, pYTK047, pYTK072, pYTK079, pYTK081 and pYTK083. Plasmid pUDE480 expressing mRuby2 was constructed from GFP dropout plasmid pUD429 with pYTK011, pYTK034 and pYTK054 using golden gate assembly as described previously (18). Similarly, pUDE481 expressing mTurquoise2 was constructed from pUD430, pYTK009, pYTK032 and pYTK053, and pUDE482 expressing Venus from pUD431, pYTK013, pYTK033 and pYTK055. Plasmids pUDR323, pUDR324, pUDR325, pUDR358, pUDR359, pUDR360, pUDR361 and pUDR362, expressing gRNAs targeting *SIT1, FAU1, Cas9, UTR2, FIR1, AIM9, YCK3* and intergenic region 550K respectively, were constructed using NEBuilder^®^ HiFi DNA Assembly Master Mix by assembling the 2 μm fragment amplified from pROS11 with primers 12230, 12235, 9457, 12805, 12806, 12807, 12808, 12809 respectively, and the plasmid backbone amplified from pROS11 with primer 6005 as described previously (3).

Plasmid pUDP045, expressing gRNA_MAL11_ and *cas9*, was constructed in a one-pot reaction by digesting pUDP004 and pUD574 using BsaI and ligating with T4 ligase. Correct assembly was verified by restriction analysis using PdmI.

### Strain construction

Yeast strains were transformed according to the high-efficiency protocol by Gietz *et al* (19). IMX1544 was constructed by transforming IMX581 with 1 μg pUDR323 and 1 μg of a repair fragment amplified from pUD481 using primers 12233 and 12234 containing an expression cassette for mTurquoise2 and 60 bp homology arms with the *FAU1* locus. IMX1555 was constructed by transforming IMX1544 with 1 μg pUDR324 and 1 μg of repair fragment amplified from pUD480 using primers 12228 and 12229 containing an expression cassette for mRuby2 and 60 bp homology arms with the *SIT1* locus. Transformants were selected on SM-AC plates, three single colony isolates were grown overnight on YPD an streaked on SM-FAC plates. Genomic DNA of a single colony was extracted, insertion of mTurquoise2 in *FAU1* was confirmed by PCR using primers 12236 and 12237, and insertion of mRuby2 in *SIT1* was confirmed by PCR using primers 12231 and 12232 followed by digestion with PvuII and XhoI digestion. IMX1557 was constructed by adding 10 µL of stationary phase culture of IMX1555 and of IMK439 in 1 mL of SM medium, incubating overnight at 30 °C and plating on SM plates with 10 g·L^-1^ clonNAT and 100 g·L^-1^ G418. IMX1585 was constructed by adding 10 µL of stationary phase culture of IMX1555 and of S288C in 1 mL of SM medium, incubating overnight at 30°C and plating on SM plates with 10 g·L^-1^ clonNAT without added uracil. All constructed strains were grown overnight in YPD and fluorescence corresponding to mRuby2 and mTurquoise2 was verified by flow cytometry.

### Cas9 mediated targeting in *S. cerevisiae* x *eubayanus* hybrid IMS0408

IMX1421, IMX1422, IMX1423 and IMX1424 were constructed by transforming IMS0408 with 1 µg pUDP045 and 1 µg of a 120 bp repair fragment constructed by annealing primers 10813 and 10814 as described previously (8). Transformants were selected on SM-AC plates, genomic DNA of 10 single colonies was extracted, but no band could be obtained when amplifying the *MAL11* locus using primer sets 1084/1470 and 1657/1148. The exact same procedure was performed without the addition of the 120 bp repair fragment. Four randomly selected colonies transformed with repair fragment were re-streaked three times on YPD agar, the plasmid was counter-selected for by plating on SM-FAC and the isolates were stocked as IMX1421, IMX1422, IMX1423 and IMX1424.

### Cas9 mediated introduction of DSBs in *S. cerevisiae* strains

DSBs were introduced by transforming yeast strains using 1 µg of purified gRNA expression plasmid and 1 µg of gel-purified double stranded repair fragment. The expression of gRNAs was done with plasmids pMEL11 to target *CAN1*, pUDR325 to target c*as9*, pUDR358 to target *UTR2*, pUDR359 to target *FIR1*, pUDR360 to target *AIM9*, pUDR361 to target *YCK3* and pUDR362 to target 550K according to Mans *et al* (3). Repair fragments containing Venus expression cassettes were PCR amplified from plasmid pUDE482 with primers with an overlap of about 20 bp with the nucleotides flanking the targeted open reading frame and purified on a 1% agarose gel (Table S5). Upon transformation, the cells were transferred to 100 mL shake flasks containing 20 mL SM-AC medium and grown until stationary phase at 30°C and 200 RPM to select cells transformed with the gRNA expression plasmid. After about 72h, 0.2 mL of these cultures was transferred to fresh SM-AC and grown under the same conditions to stationary phase to dilute any remaining untransformed cells. After about 48h, 0.2 mL of these cultures was transferred to 100 mL shake flasks containing 20 mL YPD medium and grown for about 12h under the same conditions to obtain optimal fluorescence signals.

### DNA extraction and whole genome analysis

IMX1557, IMX1585, IMX1596-IMX1635, IMS0408 and IMX1421-IMX1424 were incubated in 500 mL shake flasks containing 100 mL liquid YPD medium at 30 °C on an orbital shaker set at 200 RPM until the strains reached stationary phase with an OD_660_ between 12 and 20. Genomic DNA was isolated using the Qiagen 100/G kit (Qiagen, Hilden, Germany) according to the manufacturer’s instructions and quantified using a Qubit^®^ Fluorometer 2.0 (ThermoFisher Scientific). Between 11.5 and 54.6 µg genomic DNA was sequenced by Novogene Bioinformatics Technology Co., Ltd (Yuen Long, Hong Kong) on a HiSeq 2500 (Illumina, San Diego, CA) with 150 bp paired-end reads using TruSeq PCR-free library preparation (Illumina). For IMX1557, IMX1585 and IMX1596-IMX1635, reads were mapped onto the *S. cerevisiae* CEN.PK113-7D genome (13) using the Burrows–Wheeler Alignment tool (BWA) and further processed using SAMtools and Pilon for variant calling (20-22). Homozygous SNPs from IMX1585 were subtracted from the list of homozygous SNPs of each strain and a list of homozygous SNPs on chromosome V was compiled per strain. Based on the list of heterozygous SNPs in IMX1585, all homozygous SNPs corresponded to the nucleotide from S288C while the nucleotide from IMX1557 was lost, and regions were identified in which all contiguous heterozygous SNPs lost heterozygosity for each strain. LOH was confirmed by visualising the generated.bam files in the Integrative Genomics Viewer (IGV) software (23). Regions mapped as having lost heterozygosity correspond to regions between the first and last nucleotide which lost heterozygosity. For IMS0408 and IMX1421-IMX1424, reads where aligned to a reference genome obtained by combining the reference genome of CEN.PK113-7D (13) and the reference genome of *S. eubayanus* strain CBS12357 (24) as they are closely related to the haploid parents of IMS0408. Regions affected by loss of heterozygosity were defined as regions in which reads did not align to the *S. cerevisiae* reference chromosome VII while reads aligned to the corresponding region of the *S. eubayanus* reference chromosome VII with approximately double the normal coverage.

## RESULTS

### Targeting of a heterozygous gene in a *S. cerevisiae* x *eubayanus* hybrid

To investigate Cas9 gene editing in a genetic context with extensive heterozygosity, we targeted a heterozygous locus in an interspecific *S. cerevisiae* x *eubayanus* hybrid. The hybrid IMS0408 was constructed previously by mating a haploid *S. cerevisiae* laboratory strain and a haploid spore from the *S. eubayanus* type strain CBS 12357, resulting in an allodiploid strain with approximately 85% nucleotide identity between corresponding chromosomes of the two subgenomes (25). The *MAL11* gene encodes a membrane transporter located on chromosome VII in *S. cerevisiae*, which is absent in *S. eubayanus* CBS 12357 genome. Therefore, the *S. cerevisiae* chromosome VII could be specifically targeted using Cas9 and a gRNA targeting *MAL11*. IMS0408 was transformed with plasmid pUDP045, expressing Cas9 and a gRNA targeting *MAL11*, with and without a repair fragment with 60-bp of homology to sequences adjacent to the 5’ and 3’ ends of the coding region of *MAL11*. Normally, selection for the presence of the Cas9/gRNA expression plasmid is sufficient to obtain accurate gene editing in almost 100% of transformed cells without the need of a selection marker incorporated in the repair fragment in Saccharomyces yeast (3,8). In common laboratory strains, replacement of a sequence with a repair DNA is commonly detected by diagnostic PCR. However, in the hybrid strain IMS0408, multiple attempts failed to yield the expected fragments after transformation with the gRNA targeting *MAL11* and a repair fragment. Therefore, the genomes of four random transformants, named IMX1421 to IMX1424, were sequenced using 150 bp paired-end Illumina reads and aligned to a haploid *S. cerevisiae* x *S. eubayanus* reference genome. While reads of strain IMS0408 aligned unambiguously to the *MAL11* locus on chromosome VII of the *S. cerevisiae* sub-genome, *MAL11* DNA was absent in transformants IMX1421-IMX1424. Absence of *MAL11* was associated with loss of large regions of chromosome VII, ranging from 29 to 356 kbp (Fig. 1). Concomitantly, the corresponding regions on the *S. eubayanus* chromosome VII devoid of *MAL11* orthologue showed double sequence coverage, indicating that targeting of *MAL11* using Cas9 resulted in replacement of varying regions of the targeted chromosome by corresponding regions from the homeologous chromosome (Fig. S1). These results indicated that genome editing using Cas9 caused loss of heterozygosity rather than the intended gene editing when targeting a locus present on just one of two homologous chromosomes in a heterozygous yeast.

**Figure 1.**
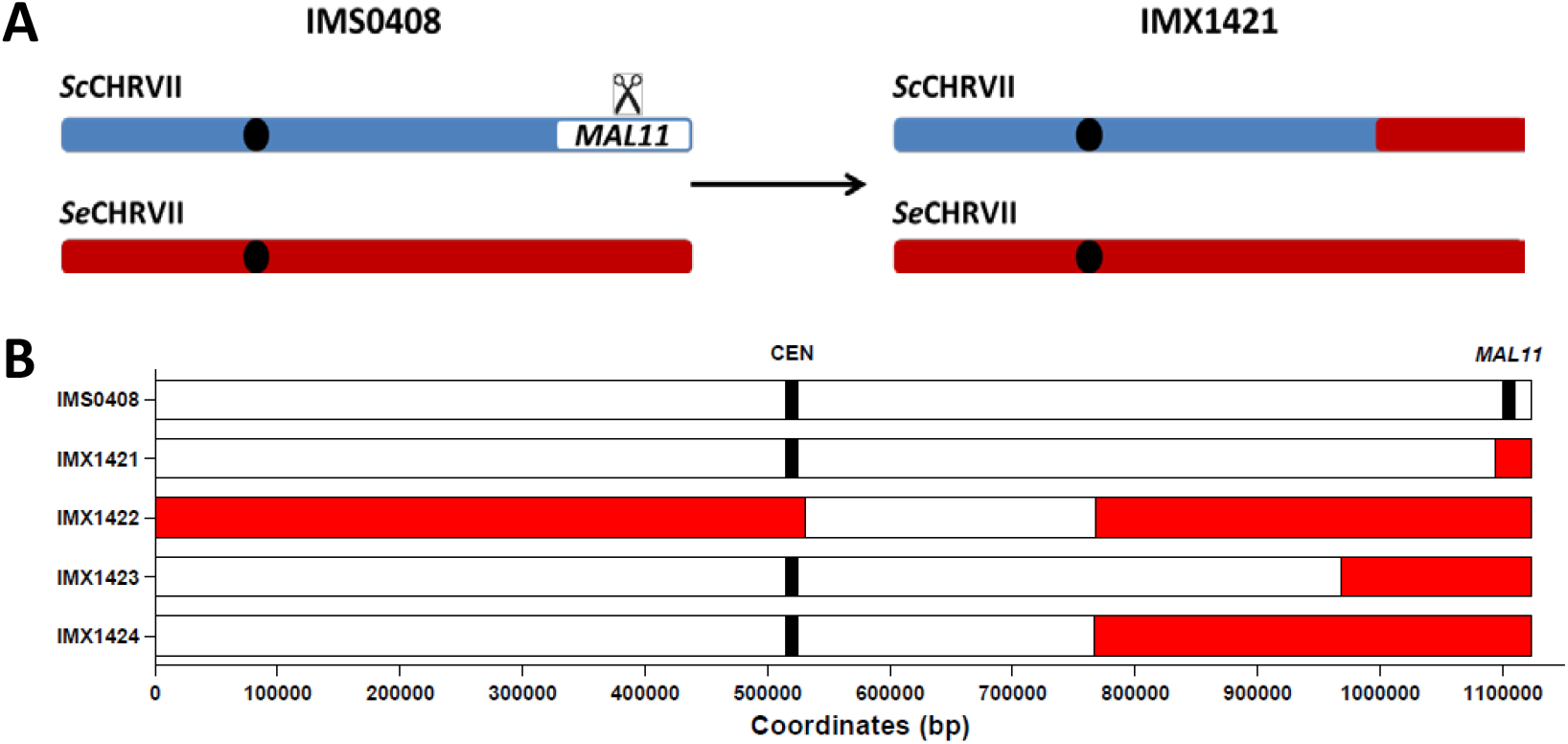
Loss of heterozygosity observed by whole genome sequencing upon Cas9-targeting of *MAL11* on the *Saccharomyces cerevisiae* derived chromosome VII in the *S. cerevisiae* x *eubayanus* hybrid IMS0408. IMS0408 was transformed with a 120 bp repair fragment with 60 bp flanks corresponding to the sequence before and after *MAL11* and with plasmid pUDP045 expressing Cas9 and a gRNA targeting the *S. cerevisiae* specific gene *MAL11* gene. Upon plating on selective medium, four randomly picked colonies were selected and sequenced using 150 bp pair-end reads and mapped against a reference genome composed of chromosome level assemblies of *S. cerevisiae* and of *S. eubayanus*. The centromere and targeted gene *MAL11* are shown at their exact coordinates, but their size is not at scale. Loss of heterozygosity is shown in red and was defined as regions in which reads did not align to the *S. cerevisiae* reference chromosome VII while reads aligned to the corresponding region on the *S. eubayanus* reference chromosome VII with approximately double the normal coverage.

### Targeting of heterozygous loci in a mostly homozygous diploid *S. cerevisiae* strain

To investigate if the observed lack of efficient gene editing was specific to this highly heterozygous *S. cerevisiae* x *eubayanus* hybrid, we systematically investigated the impact of target-sequence heterozygosity on the efficiency of gene editing in *S. cerevisiae* strains. To this end, DSBs were introduced at homozygous and heterozygous loci on chromosome V of yeast strains that carried a Cas9 expression cassette integrated at the *CAN1* locus. Plasmid-based gRNA expression was performed as described previously (3). Use of a repair fragment expressing the fluorescent protein Venus enabled analysis of editing efficiency by flow cytometry (18). To verify functional Cas9 and gRNA expression, the *Δcan1::cas9* locus was first targeted in the haploid *S. cerevisiae* strain IMX1555, resulting in integration of the repair fragment in 98.3±1.3% of cells (Table S1). Subsequently, the homozygous alleles of *AIM9* and *YCK3* were targeted in the congenic diploid *S. cerevisiae* strain IMX1557, resulting in integration of the repair fragment in 98.6±0.8% and 99.2±0.4% of cells, respectively (Fig. 2A). In contrast, when individually editing each allele of the heterozygous *CAN1*/*Δcan1::cas9* locus in the diploid strain IMX1557, the repair fragment was integrated in only 4.4±2.5% of cells when targeting the *Δcan1::cas9* allele, and 0.9±0.6% of the cells when targeting the *CAN1* allele (Fig. 2A). These results indicated that gene editing efficiencies were up to 99-fold lower for heterozygous target loci than for homozygous target loci (p<10^-4^). Since IMX1557 was homozygous in most of its genome, except the targeted locus, the introduction of a DSB in only one of two homologous chromosomes rather than genome heterozygosity itself, impeded accurate and efficient gene editing using Cas9.

**Figure 2.**
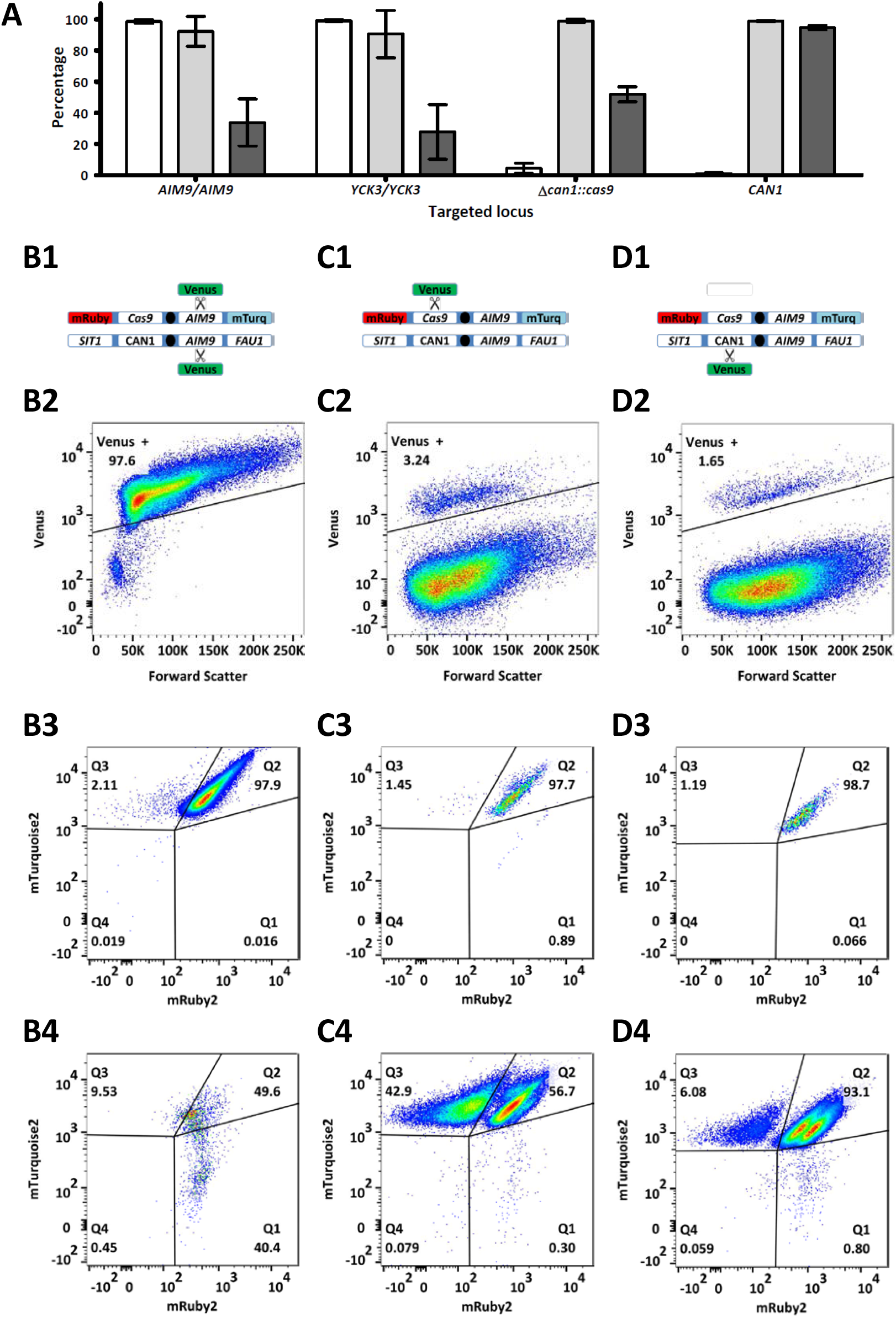
Cas9-mediated gene editing of homozygous and heterozygous loci on chromosome V of *S. cerevisiae*. (A) Average fluorescence of cell populations in which the homozygous *AIM9* and *YCK3* alleles and the heterozygous *Cas9* and *CAN1* alleles were targeted in the diploid strain IMX1557. The percentage of cells expressing Venus (white), the percentages of cells expressing both mTurquoise2 and mRuby2 in the Venus positive (light grey) and Venus negative cells (dark grey) are shown. For each target, averages and standard deviation for biological triplicates are shown. **(B,C and D) Fluorescence profiles obtained when targeting *AIM9*, Cas9 and *CAN1* in IMX1557.** (row 1) Schematic representation of both copies of chromosome V in IMX1557, with the alleles at the *SIT1, CAN1, AIM9* and *FAU1* loci and scissors indicating Cas9 targeting. While one chromosome copy has the wildtype alleles for all loci, the other copy has mRuby2 integrated in *SIT1*, Cas9 integrated in *CAN1* and mTurquoise2 integrated in *FAU1*. (rows 2, 3 and 4) Flow cytometry profiles of targeted cells. Each gene was targeted in three biological replicates and flow cytometric data for a representative replicate is shown. After transformation, 100,000 cells were analysed by flow cytometry and single cells were selected based on a FSC-A/FSC-H plot to avoid multicellular aggregates. For each replicate, at least 75,000 single cells remained and the fluorescence corresponding to Venus was used to determine gene-editing efficiency (row 2). For each gene, the fluorescence corresponding to mRuby2 and mTurquoise2 is plotted for the cells with (row 3) and without (row 4) expression of Venus. Fluorescence results for all samples are provided in Table S1.

In order to investigate if Cas9 gene editing resulted in loss of heterozygosity, as observed in the hybrid IMS0408, the presence of both chromosome arms of the targeted chromosome homolog was monitored by flow cytometry. IMX1557 expressed the fluorophores mRuby2 and mTurquoise2 from the *SIT1* and *FAU1* loci of the chromosome V copy harbouring the *Δcan1::cas9* allele, but not from the non-modified homologous chromosome (Fig. B1-D1). Loss of the left and right arms of the copy of chromosome V harbouring *Δcan1::cas9* could therefore be monitored by measuring fluorescence corresponding to respectively mRuby2 and mTurquoise2 (18). After expressing a gRNA targeting the cas9 cassette, 99.4±0.3% of cells still expressed mTurquoise2. However, while 99.5±0.7% of the correctly gene-edited cells still expressed mRuby2, 47.6±2.7% of the Cas9-targeted cells that did not integrate the repair fragment had lost mRuby2 fluorescence (Fig. 2A). These results indicated that targeting of a heterozygous locus resulted in loss of sequences on the targeted chromosome arm, but did not affected the opposite chromosome arm. Similarly, after targeting the *CAN1* allele of the same locus, two distinct subpopulations were discernible in cells that had not integrated the repair fragment (Fig. 2D). The two-fold difference in mRuby2 fluorescence between these two subpopulations could reflect duplication of mRuby2. Loss of mRuby2 fluorescence upon transformation with a gRNA targeting *Δcan1::cas9* and doubling of mRuby2 fluorescence when targeting *CAN1* were also observed in the absence of a co-transformed repair fragment (Table S1). This indicates that introduction of a DSB at a heterozygous locus caused loss of heterozygosity (LOH) through replacement of a targeted chromosome segment by duplication of the corresponding segment from its homologous chromosome, as was observed when targeting *MAL11* in the *S. cerevisiae* x *eubayanus* hybrid IMS0408.

### Elucidation of mutations caused by Cas9-targeting using whole genome sequencing

Chromosome-arm LOH has previously been reported upon introduction of a DSB in one of two homologous chromosomes, but was considered rare and has not been described as disruptive to gene-editing approaches (9,26,27). To investigate the extent and nature of the LOH caused by Cas9-editing of heterozygous loci, a strain with approximately four heterozygous SNPs or INDELs per kbp was generated by mating IMX1555 (CEN.PK genetic background, expressing Cas9, mRuby2 and mTurquoise2 from chromosome V) with S288C (Table S6). LOH could be monitored at the chromosome arm level by flow cytometry and at the nucleotide level by whole-genome sequencing. By using PAM sequences absent in S288C, we specifically targeted the CEN.PK-derived chromosome V, which carried expression cassettes for mRuby2 and mTurquoise2 on its left and right arms, respectively, at the *CAN1, UTR2, FIR1, AIM9* and *YCK3* loci and at intergenic coordinate 549603, referred to as 550K. Upon targeting of the *CAN1* and *UTR2* loci, mRuby2 fluorescence was lost in 46.7±2.4 and 11.2±0.2% of cells, respectively, while mTurquoise2 fluorescence was unaffected in at least 99.6±0.2% of the cells (Fig. 3A). Targeting of the *FIR1, AIM9, YCK3* or 550K loci caused loss of mTurquoise2 fluorescence in 12.2±0.4, 13.6±0.1, 12.7±0.2 and 43.6±0.3% of cells, respectively, while mRuby2 fluorescence was conserved in at least 98.1±0.5 % of cells (Fig. 3A). As the centromere is located between *UTR2* and *FIR1*, these results confirm that, for all investigated loci, a large fraction of cells lost the targeted chromosome arm. Fluorescence-assisted cell sorting (FACS) was subsequently used to isolate 10 single cells each from the following populations: *UTR2*-targeted cells with mRuby2 fluorescence (IMX1606-IMX1615), *UTR2*-targeted cells without mRuby2 fluorescence (IMX1596-IMX1605), *FIR1*-targeted cells with mTurquoise2 fluorescence (IMX1626-IMX1635), and *FIR1*-targeted cells without mTurquoise2 fluorescence (IMX1616-IMX1625). Whole-genome sequencing and alignment of reads to the CEN.PK113-7D genome sequence (13) revealed LOH of the targeted locus in all 40 isolates (Fig. 3B). In cell lines that did not lose a fluorophore, LOH was local, affecting regions ranging from 3 to 17,495 nucleotides for *UTR2*-targeted cells and regions ranging from 1 to 11,900 nucleotides for *FIR1*-targeted cells, corresponding to up to 79 mutations (Fig. 3C and Table S2). In isolates that did lose a fluorophore, LOH affected the chromosome arm harbouring the targeted locus, affecting 79,859 to 110,289 nucleotides for *UTR2*-targeted cells and 359,841 to 362,790 nucleotides for *FIR1*-targeted cells, corresponding to up to 1,697 mutations (Fig. 3C and Table S2). Absence of newly introduced SNPs at targeted loci indicated that repair of DSBs did not involve non-homologous end joining (28).

**Figure 3.**
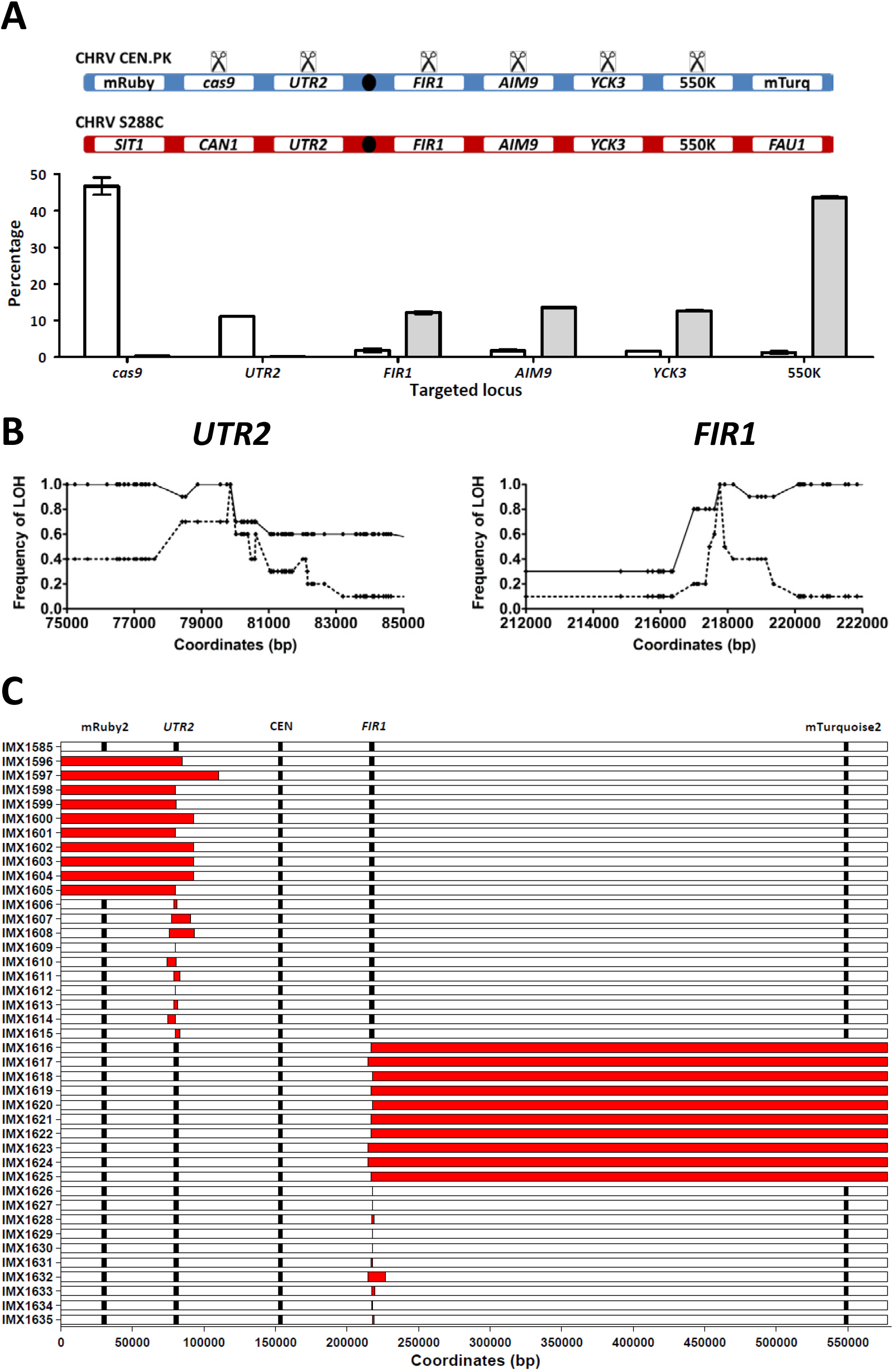
Loss of heterozygosity caused by Cas9-mediated gene editing at heterozygous loci in the heterozygous *S. cerevisiae* diploid IMX1585. Mating of the haploid *S. cerevisiae* strains IMX1555 (CEN.PK-derived) and S288C yielded the heterozygous diploid strain IMX1585 (ca. 4 heterozygous nucleotides per kbp on chromosome V). The CEN.PK-derived chromosome harbours the fluorophores mRuby2 and mTurquoise2, enabling detection of the loss of each arm of the CEN.PK-derived chromosome V by flow cytometry. DSBs were introduced specifically in the CEN.PK–derived chromosome and loss of heterozygosity was monitored at the population level using flow cytometry and in single cell isolates by whole genome sequencing. **(A) Population-level loss of heterozygosity after targeting *cas9, UTR2, FIR1, AIM9 YCK3* and 550K in IMX1585.** In a schematic representation of the CEN.PK-derived and S288C-derived chromosome V, targeted loci are indicated by scissors, the fluorophores cassettes by their respective fluorescent colour and the centromere by a black oval. In the graph, the percentage of cells having lost mRuby2 fluorescence (white) and mTurquoise2 (red) is shown for each targeted locus. Averages and standard deviations were calculated from biological triplicates. **(B) Loss of heterozygosity at the nucleotide level in single isolates obtained by targeting *UTR2* and FIR1 in IMX1585.** For each targeted locus (indicated by scissors), the frequency of LOH is shown for 10 isolates with intact fluorescence (dashed line, IMX1606-IMX1615 and IMX1626-IMX1635) and 10 isolates having lost a fluorophore (continuous line, IMX1596-IMX1605 and IMX1616-IMX1625. **(C) Overview of loss of heterozygosity across chromosome V in isolates in which *UTR2* and *FIR1* were targeted using Cas9.** The non-targeted strain (IMX1585), *UTR2*-targeted isolates with fluorescence of mRuby2 and mTurquoise2 (IMX1606-IMX1615), *UTR2*-targeted isolates which lost mRuby2 fluorescence (IMX1596-IMX1605), *FIR1*-targeted isolates with fluorescence of mRuby2 and mTurquoise2 (IMX1626-IMX1635) and *FIR1*-targeted cells which lost mTurquoise2 fluorescence (IMX1616-IMX1625) were sequenced using 150 bp paired-end reads and mapped against the CEN.PK113-7D genome. The fluorophores mRuby2 and mTurquoise2, the targeted genes *UTR2* and *FIR1* and the centromere are shown at their exact coordinates, but their size is not at scale. Loss of heterozygosity was defined as regions in which nucleotides which were heterozygous in IMX1585 were no longer heterozygous in the isolate (in red). Exact coordinates are provided in Table S2.

### Identification of repair patterns corresponding to homology-directed repair

We conclude that introduction of a DSB at a heterozygous locus results in low gene-editing efficiencies due to a competing repair mechanism that causes local or chromosome-arm LOH. While repair using homologous chromosomes typically relies on BIR, HR or HDR in eukaryotes (29), the observed local LOH is consistent with HDR (Fig. 4A) (10-12). Indeed, strains IMX1606, IMX1608 and IMX1613 showed patterns of alternating homozygous and heterozygous sequences around the targeted locus consistent with the heteroduplex resolution step characteristic for HDR (Fig. 4A and Table S2). Although previous studies attributed chromosome-arm LOH to BIR or HR (9,26), occurrence of similar mosaic structures in strains with chromosome-arm LOH (strains IMX1605 and IMX1619, Table S2) indicated that HDR was also responsible for chromosome-arm LOH. While BIR or HR do not cause mosaic LOH, chromosome-arm LOH is not a commonly-recognized result of HDR (Fig. 4A) (10-12). However, we propose a repair mechanism that involves HDR of one of the targeted chromatids at the 2n stage of the cell cycle (Fig. 4B), which is consistent with all phenotypes and genotypes encountered in this study as well as in previous studies involving hemizygous introduction of DSBs (9,26,27,30).

**Figure 4.**
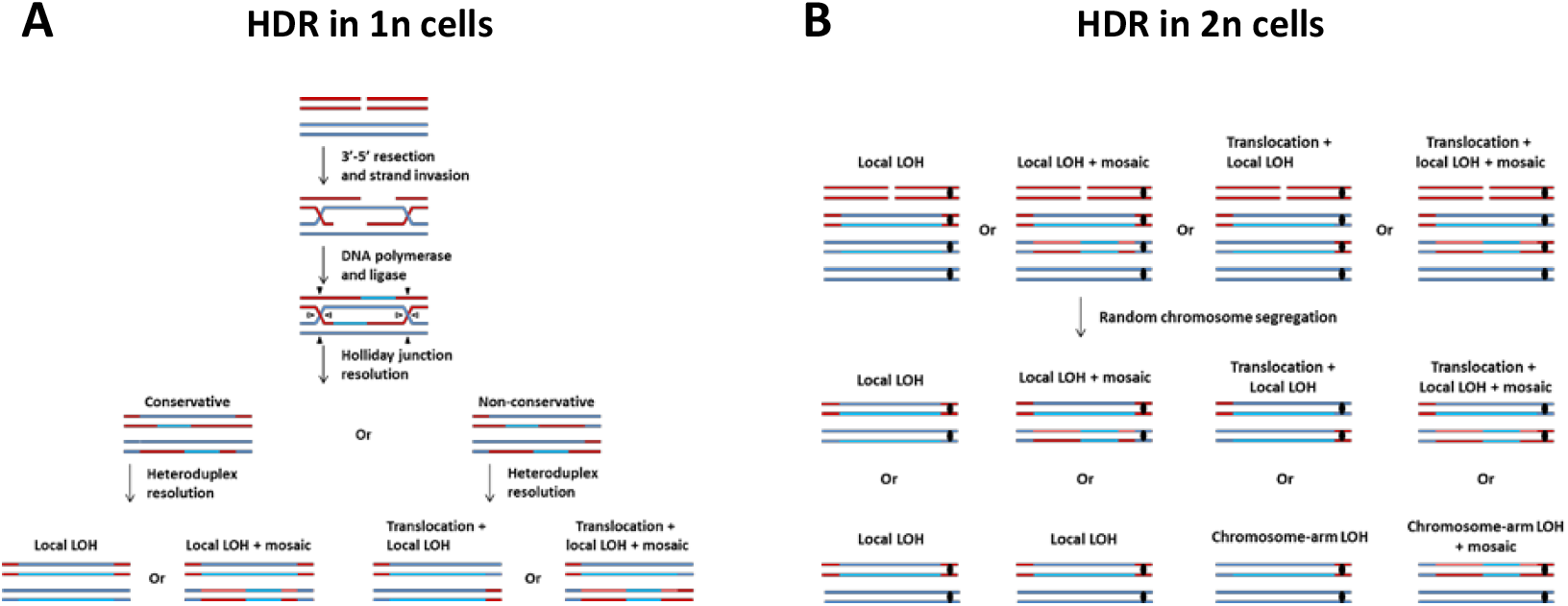
Proposed mechanism for Cas9-mediated loss of heterozygosity based on homology-directed repair (HDR) between homologous chromosomes. (A) Possible outcomes of HDR in cells with one chromosome complement (1n). (B) Possible outcomes of HDR in cells with two chromosome complements (2n). The targeted chromosome (red), its homolog (blue) and the centromere are indicated (black, where relevant). Newly synthesized DNA is shown in a lighter shade. During heteroduplex resolution, the strand with the targeted NGG PAM sequence is always discarded due to Cas9 activity. For 2n HDR, HDR occurs between one chromatid of the targeted and one chromatid of the non-targeted chromosome, as in 1n HDR. The chromatids subsequently segregate according to their centromere pairing, with one red and one blue centromere in each daughter cell. Cells receiving the unrepaired red chromosome die. As indicated in the figure, HDR in 2n cells could yield local as well as chromosome-arm LOH, both with and without mosaic structures.

## DISCUSSION

The efficiency of gene editing using Cas9 can decrease by almost two orders of magnitude when targeting only one of two homologous chromosomes due to a competing repair mechanism causing either local or chromosome-arm scale LOH. Contrarily to previously identified side effects of cas9-mediated gene editing, the observed LOH consisted not only of loss of genetic material from the targeted chromosome (31), but also of replacement of the affected sequence by an additional copy of sequence homologous to the targeted site. While such LOH upon introduction of a hemizygous DSB has been observed in the yeasts *S. cerevisiae* and *Candida albicans* (9,26), this study demonstrates that repair by LOH is not only possible, but occurs at rates which impede gene editing approaches based on integration of repair fragments. This phenomenon is likely to contribute to a lesser genome accessibility of heterozygous yeasts relative to laboratory strains, which tend to be haploid or homozygous. Therefore, these results are likely to affect the genome editing of hybrids, industrial yeasts and natural isolates due to their frequent heterozygosity (32), and should be used to update guidelines for designing gene editing strategies. We strongly recommend to design gRNAs targeting homozygous nucleotides stretches when targeting heterozygous genomes. When allele-specific gene editing is required, we recommend the use of repair fragments with integration markers such as the Venus fluorophore in this study, since accurate gene editing is not impossible, simply inefficient. When the use of a marker is not permissible, extensive screening of transformants for correct gene editing may be required.

While the HDR machinery is well conserved in eukaryotes (11,12), further research is required to determine if LOH occurs at similar rates in eukaryotes other than *S. cerevisiae*, and if it impedes gene editing. While DSB-mediated LOH was observed in *S. cerevisiae, C. albicans, Drosophilia melanogaster* and *Mus musculus* (9,26,27,30), relative contributions of HR, HDR and NHEJ to DSB repair vary across species. However, since integration of a repair fragment and repair by LOH both involve HDR (33,34), targeting heterozygous loci likely causes gene-editing efficiencies and off-target mutations in other eukaryotes as well, regardless of the efficiency of NHEJ and HR.

Targeting of heterozygous loci is common in gene editing, for example during allele propagation of gene drives and disease allele correction in human gene therapy (33,34). Although gene drives are based on LOH by HDR (34), the extent of LOH beyond the targeted locus has not been systematically studied but could, by analogy with the present study, potentially affect entire chromosome arms. Allele-specific gene editing generally aims at repair by HDR using a co-transformed repair fragment instead of a homologous chromosome. Reports of LOH after targeting a heterozygous allele in human embryos despite availability of an adequate repair fragment, are consistent with Cas9-induced LOH extending beyond the targeted locus, as described here (33). While, in the human-embryo study, repair by LOH was perceived as a success, the reported role of LOH in cancer development (35) indicates that large-scale LOH can have important phenotypic repercussions. Therefore we recommend avoiding allele-specific gene editing when possible until further research determines if it is a risk in other eukaryotes. Based on the proposed HDR mechanism for CRISPR/Cas9-mediated LOH (Fig. 4B), the risk of LOH can be mitigated by designing gRNAs that cut all alleles of heterozygous loci, even if only a single allele needs to be edited. Eventually, CRISPR-Cas9 editing could become safer by favouring DSB-independent gene-editing methods such as guided nickases and base-editing strategies for preventing or reducing the incidence of LOH (36-39).

## ACCESSION NUMBERS

The sequencing data were deposited at NCBI (https://www.ncbi.nlm.nih.gov/) under the Bioproject PRJNA471787.

## SUPPLEMENTARY DATA

Supplementary Data are available at NAR online.

## ACKNOWLEDGEMENT

ARGdV conceived the study and designed the experiments. ARGdV and LGFC performed plasmid and strain construction. ARGdV, LGFC, PdlTC and JtH performed the experimental work. ARGdV and MvdB performed bioinformatics analysis. ARGdV, JTP and JMGD supervised the study and wrote the manuscript. All authors read and approved the final manuscript.

We thank Liset Jansen for drawing our attention to the difficulty to edit a heterozygous gene, Robert Mans for his expertise with gene editing in *Saccharomyces cerevisiae*, Melanie Wijsman for constructing and Pascale Daran-Lapujade for sharing plasmids pUDE480, pUDE481 and pUDE482, Sai T. Reddy for his insights in the potential impact for human gene therapy and Nick Brouwers, Alex Salazar, Xavier D. V. Hakkaart, Ioannis Papapetridis, Niels G.A. Kuijpers, Jan-Maarten Geertman and Thomas Abeel for their critical input.

## FUNDING

This work was supported by the BE-Basic R&D Program (http://www.be-basic.org/), which was granted an FES subsidy from the Dutch Ministry of Economic Affairs, Agriculture and Innovation (EL&I).

## CONFLICT OF INTEREST

The authors declare no conflict of interest.

